# A Functional Exodermal Suberin is Key for Plant Nutrition and Growth in Potato

**DOI:** 10.1101/2023.09.14.557788

**Authors:** Dolors Company-Arumí, Carlota Montells, Mònica Iglesias, Eva Marguí, Dolors Verdaguer, Katarina Vogel-Mikus, Mitja Kelemen, Mercè Figueras, Enriqueta Anticó, Olga Serra

## Abstract

Angiosperm roots, except in Arabidopsis, have both endodermis and exodermis, which regulate radial water and solute movement through lignin and suberin deposition. While endodermal suberin in Arabidopsis acts as a barrier to water and solute uptake and backflow, its implications in other angiosperms with both layers and the role of exodermal suberin remain unclear. We examined potato roots (*Solanum tuberosum*) and found that exodermis lacks the typical Casparian strip but forms an outer lignin cap, and quickly suberizes near the root tip. In contrast, a few endodermal cells, with Casparian strip, start suberizing much later. The continuous early exodermal suberization covering the root underlines its potential role in mineral nutrient radial movement. To demonstrate it, we used plants downregulating the suberin biosynthetic gene *CYP86A33*, which had the root suberin reduced in a 61%. Phenotypic analyses of the suberin-deficient mutant showed altered mineral nutrient concentration, slightly reduced water content and compromised growth. Micro-PIXE analyses identified the distribution of elements within the roots and highlighted anatomical compartments defined by apoplastic barriers. These findings advance our understanding of nutrient radial transport, demonstrate exodermal suberin as a bidirectional and selective barrier to element movement, and underscore its importance in nutrient homeostasis and plant growth.

## INTRODUCTION

The young roots of virtually all vascular plants develop an endodermis, a single-layer tissue surrounding the stele that forms an inner boundary of the cortex (Geldner, 2013). Additionally, approximately 90% of angiosperms differentiate a specialized hypodermis in their roots, the exodermis, structurally similar to the endodermis and usually uniseriate, which forms the outer boundary of the cortex separating it from the epidermis and surrounding medium (Perumalla et al., 1990b; Peterson and Perumalla, 1990; Peterson, 1989). Consequently, most roots of flowering plants, including crops, have a cortex cylinder enclosed by endodermis and exodermis (Hose et al., 2001; Peterson, 1989). In contrast, no exodermis has been reported for gymnosperm roots, and for seedless early vascular plants the exodermis has only been identified in the roots of the lycophyte genus *Selaginella* (Damus et al., 1997). The rhizomes of angiosperms also contain an exodermis (Perumalla et al., 1990a).

The endodermis and exodermis originate from the ground meristem of the root apical meristem. Once formed, their cells differentiate while accumulating lignin, suberin, and polysaccharides in their cell walls, thus defining functional apoplastic diffusion barriers. Differentiation state I corresponds to the formation of the Casparian strip, a lignified belt of the radial and transverse primary walls that is structurally linked to the plasma membrane (Geldner, 2013), restricted to the mid-region in the endodermis, and often occupies the whole of the radial and transverse walls in the exodermis (Hose et al., 2001). State II corresponds to the formation of suberin lamellae as an inner secondary layer encapsulating the protoplast. While endodermal differentiation can be completed with the formation of Casparian strip or progress to the formation of suberin lamellae, virtually all exodermal cells differentiate into both Casparian strip and suberin lamellae, indicating that exodermis invariably progresses to state II (Perumalla et al., 1990b; Peterson, 1989). These cells may later progress to state III by developing a tertiary polysaccharide cell wall, although it is still unclear whether this is a general feature of the endodermis and exodermis. Both the endodermis and exodermis include passage cells located at xylem poles, which form Casparian strips but delay the entry to suberization and later progression to the state III.

The endodermis and exodermis function as dynamic barriers to protect against drought, ion toxicity, radial oxygen loss (under root anoxia) and intruders (*see for review* Hose et al., 2001; Geldner, 2013; Enstone et al., 2003; Soukup and Tylová, 2018). For example, apoplastic deposits aid in explaining the capacity of young roots to selectively absorb water and nutrients and radially transfer them to the stele, where they can further reach the entire plant. Moreover, by accelerating or delaying the differentiation of the endodermis and exodermis layers, roots are expected to modulate their radial transport capacity in response to changing physiological and environmental conditions (Hose et al., 2001; Shukla and Barberon, 2021). In this respect, the significance of the Casparian strip and suberin lamellae has only been revealed for the endodermis through genetic studies, while evidence for the role of the exodermal apoplastic barrier is almost nonexistent. In the first differentiation stage, the Casparian strips form a network that blocks the apoplastic radial movement of water and nutrients, which, to cross the endodermis, will need to be uptaken by influx carriers at the endodermal plasma membrane through the trans-cellular pathway. Once in the endodermal protoplast, the path to the stele would need to go through endodermal efflux carriers (coupled trans-cellular pathway) or through the plasmodesmata (symplastic pathway) (Barberon and Geldner, 2014). Suberin lamellae would block the trans-cellular pathway by depositing between the plasma membrane and hydrophilic cell wall, thus obstructing the access of water and nutrients to aquaporins and influx and efflux carriers, respectively. The role of suberin in limiting mineral element and water movement in a selective and bidirectional manner has been recently established for endodermis by genetically reducing or depleting the root suberin in Arabidopsis (Wang et al., 2019; Ranathunge and Schreiber, 2011; Calvo-Polanco et al., 2021; Li et al., 2017; Barberon et al., 2016; Wang et al., 2020). In contrast, despite the wide occurrence of the exodermis in most angiosperms, the role of exodermal suberin is still unknown, probably because Arabidopsis roots develop a single-layered cortex and no exodermis. However, genetic and functional evidences are needed to disentangle the role of the exodermis in root function.

Much of the studies focused on roots containing both endodermal and exodermal layers have been carried out in the roots of important cereal crops such as barley, maize or rice or, most recently, tomato (*see i.e.* Kreszies et al., 2018; Kajala et al., 2021; Líška et al., 2016; Shiono et al., 2022; Namyslov et al., 2020), which are seed-propagated and the root system is initiated embryonically (seeds). However, studies on roots from root or tuber crops that are commonly vegetatively propagated are neglected despite their potential to contribute to food security in the future (Khan et al., 2016). Potato (*Solanum tuberosum*), the world’s most important non-cereal food crop, is propagated from tubers and its root system is composed of adventitious roots (Joshi and Ginzberg, 2021).

To gain some knowledge of exodermal suberization, we identified the suberization pattern of potato adventitious roots. The exodermis is the first layer that quickly suberizes, forming a complete suberized cylinder close to the root tip. In contrast, endodermis suberization is much delayed and occurs in particular cells in regions far from the root tip. To learn about the role of exodermal suberin in plant nutrition, we characterized the root suberin of the *CYP86A33*-RNAi, resulting in impaired suberin which amounted for 40% of the wild type. Using this suberin-deficient mutant we demonstrated the role of exodermal suberin as a selective bidirectional barrier to nutrients, and mapped the root mineral element distribution to localize the ion-specific accumulation in anatomical compartments. Biomass and water content measurements allow for the assessment of the impact of exodermal suberin deficiency on plant growth and water retention.

## RESULTS

### Potato root apoplastic diffusion barriers

We first aimed to describe the anatomy and localization of the apoplastic barriers of the primary adventitious potato roots emerging from the stem of *Solanum tuberosum* cv. Desirée plants grown in hydroponics. To detect lignin and suberin, we stained the roots using basic fuchsin and Nile red, respectively, and added calcofluor white to stain the polysaccharides from the cell walls. The stained roots were observed using confocal microscopy.

We detected a basic fuchsin signal of lignin in both the endodermis and exodermis cell layers in root regions below 2 cm from the root tip (**Figure 1**). In the endodermis, we observed the lignin signal deposited in the typical Casparian strip pattern (**Figure 1A**, white arrowheads), which formed a longitudinal continuous network surrounding the vascular cylinder (**Figure 1A**, endodermis plane). In the exodermis, we also observed lignification but did not form the typical Casparian strips. Instead, exodermal lignification was displaced to the external corners of the outer tangential cell wall and extended inward to the radial exodermal cell walls and outward to the radial epidermal cell walls (**Figure 1A**; yellow arrowheads). In the longitudinal plane, this lignification pattern creates a continuous lignified network involving the exodermal and epidermal cells (**Figure 1A**, exodermis and epidermis plane), similar to that created for the Casparian strip in the endodermis. Therefore, we could expect that lignification in both exodermal/epidermal cells and endodermal Casparian strips might contribute to creating the apoplastic diffusion barriers to the radial transport of water and solutes.

**Figure 1.**
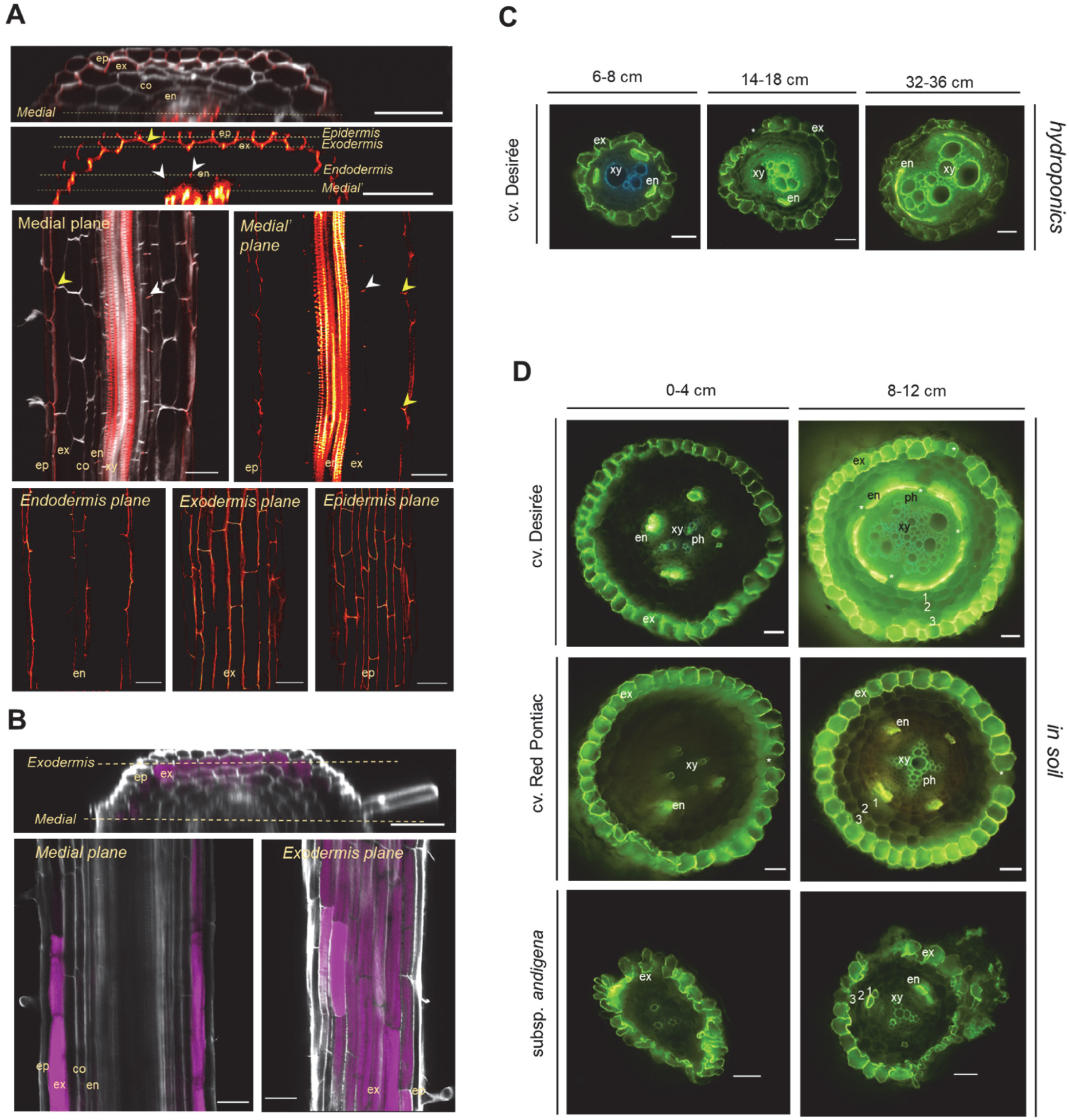
Apoplastic barriers in adventitious potato roots. (A-C) Root anatomy of *S. tuberosum* cv. Desirée grown in hydroponics. (A) Root segment at 2 cm from the root tip stained for lignin (red, basic fuchsin) and polysaccharides (white, calcofluor white) and visualized by confocal microscopy. Orthogonal view of z-scan series and single longitudinal views are shown. To better visualize the lignin, the Nile red signal is shown as a fire hot gradient, where yellow pixels are those with higher signal intensities. White arrowheads indicate the lignified Casparian strip and yellow arrowheads indicate exodermis radial cell wall lignification. (B) Root segment at 2-4 cm from the root tip stained for suberin (magenta, Nile red) and polysaccharides (white, calcofluor white) and visualized by confocal microscopy. Orthogonal view of z-scan series and single longitudinal views are shown. (C-D) Root free-hand cross-sections stained for suberin (green, fluorol yellow) and observed under the epifluorescence microscope. Representative images from sections obtained at different distances from the root tip. (C) Roots grown in hydroponics. (D) Roots grown in soil of different potato *S. tuberosum subsp. tuberosum* commercial varieties (cv.) and a wild relative *S. tuberosum subsp. andigena* grown in soil. The numbers indicate the individual cortex layers. co, cortex; en, endodermis; ep, epidermis; ex, exodermis; xy, xylem; ph, phloem. Asterisks indicate single unsuberized cells in the exodermal and endodermal layers corresponding to passage cells. The scale bars correspond to 50 μm.

To detect suberin, we first used Nile red staining. In root regions below 2 cm from the root tip, suberin was already observed in the exodermis (**Figure 1B**). The suberin signal in the exodermis was homogeneously distributed throughout the perimeter of the exodermal cell wall. We could not detect suberin in the endodermis in any of the specimens observed using samples up to 10-cm from the root tip. To confront these observations and determine whether the endodermis of potato roots grown in hydroponics was actually unsuberized, we stained free-hand root cross sections with fluorol yellow, a fluorescent dye commonly used to stain suberin. Again, suberin was clearly detected in the exodermis cells forming a continuous suberized layer already in the first 2 cm from the root tip (**Figure 1C**), except for some cells that remain unsuberized which may correspond to passage cells. Endodermal suberization was observed in particular cells in the root regions 6–11 cm from the root tip (**Figure 1C**). For roots grown in hydroponics for longer periods (seven weeks instead of three weeks), we observed an increased number of endodermal cells that deposited suberin, being much more prominent at the base of the root (approximately 32-36 cm), where vascular secondary growth was evident (**Figure 1C**).

Since exodermal suberization is described to be very reactive to environmental conditions, we wondered whether this early and intense suberization of exodermis was because the roots were growing in hydroponics or was an intrinsic characteristic of potato roots. To answer this, we analyzed the suberized layers of adventitious potato roots grown in soil using the same variety tested previously in hydroponics (cv. Desirée). Additionally, to determine the extent to which exodermal suberization may be a common characteristic in potato, we also included another *S. tuberosum* subsp*. tuberosum* cultivar, cv. Red Pontiac, and the wild relative *S. tuberosum* subsp*. andigena*. In the youngest root regions (up to 4 cm from the root tip), we observed an early exodermal suberization, which formed a continuous layer (**Figure 1D**) punctually interrupted by unsuberized passage cells located at the xylem poles (**Figure 1D**, asterisks). In contrast, in these youngest regions, any or only individual endodermal cells were suberized, usually located at the phloem pole. At mature stages (8-12 cm from the root tip), endodermis suberization was more prominent than in younger root regions, although the number of suberized cells was clearly different between the varieties. Whereas in subsp. *tuberosum* cv. Red Pontiac and subsp. *andigena* suberized endodermal cells were still restricted at the phloem pole, in subsp. *tuberosum* cv. Desirée suberization progressed to neighboring endodermal cells, forming an almost continuous suberized layer, except for some endodermal passage cells (**Figure 1D**, asterisks). Regarding the growing conditions, our data indicated that endodermal suberization is triggered earlier in soil than in hydroponic conditions (compare **Figure 1C** and **Figure 1D**).

Overall, our observations indicated that in potato roots, grown in soil and hydroponics, the exodermis is the main and often solely suberized layer in younger roots, covering almost its entire length but the root tip (**Figure 1C-D**). Suberization in the endodermis occurs much later in development, in more mature root regions, and in hydroponics, this suberization is further delayed (**Figure 1C-D**), suggesting plasticity of endodermal suberization in response to root environmental growth conditions, as observed also for Arabidopsis (Barberon et al., 2016). Hence, exodermal suberization in potato roots is the suberized cell layer that potentially restricts water and mineral element movement and contributes to nutrient homeostasis.

### The root of *CYP86A33*-RNAi line is deficient in suberin

To study the importance of suberin accumulated in exodermis for plant nutrition, we selected the *CYP86A33*-RNAi potato plant (*subsp. tuberosum* cv. Desirée background) for its expected reduction in suberin deposition in roots, based on the 60% reduction observed in the potato tuber periderm (Serra et al., 2009). Hydroponics appeared to be an exceptional model because in these conditions, the suberin deposited in the primary roots was extensively found in exodermal cell walls, while endodermis remained practically unsuberized (**Figure 1C-D**).

We first tested the *CYP86A33* gene downregulation in the roots of two *CYP86A33*-RNAi lines (22 and 39). The RT-qPCR analysis demonstrated a residual accumulation of *CYP86A33* transcript (14 and 12% respectively of that of wild type) (**Supplemental Figure 1**) and so these lines were used for further analyses.

The effect of *CYP86A33* downregulation on root suberin was first analyzed histologically in plants grown in hydroponics for three weeks. Fluorol yellow staining of *CYP86A33*-RNAi root cross-sections showed a weaker signal in the exodermal layer, as well as in individual suberized endodermal cells, than in wild-type roots (**Figure 2**) using the same epifluorescence microscopy parameters. In the exodermis, only individual cells remained unsuberized, which corresponded to passage cells. The analysis again confirmed that under such conditions the exodermis was heavily suberized, forming a continuous suberized cylinder along the root, whereas suberin in the endodermis was restricted to individual cells.

**Figure 2.**
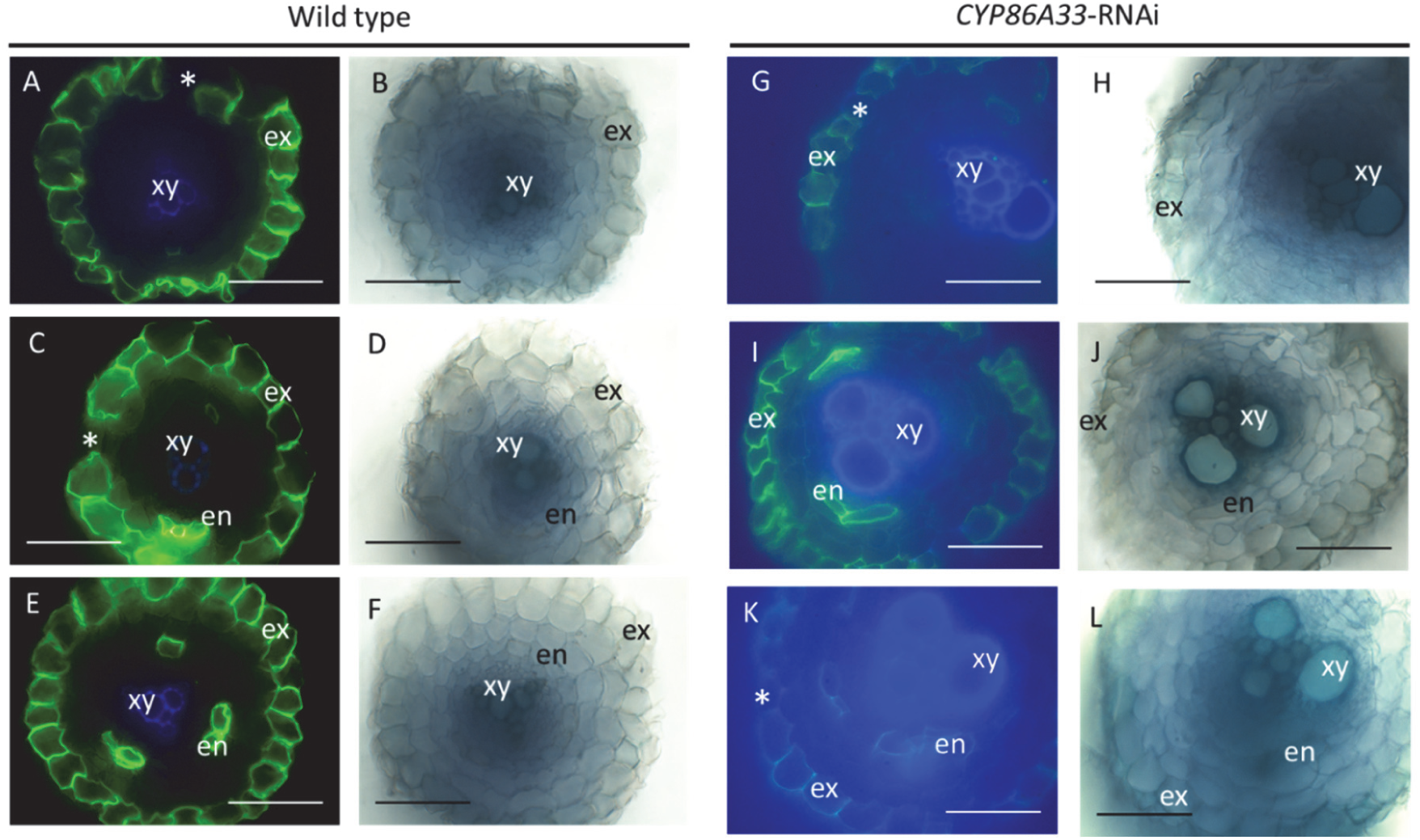
Effect of *CYP86A33* downregulation in suberin deposits by histological analyses. Free-hand root cross-sections stained with fluorol yellow (green, suberin) obtained from wild type and *CYP86A33*-RNAi potato (cv. Desirée) plants grown in hydroponics. The epifluorescence microscopy micrographs (A, C, E, G, I, K) were observed under UV filter in which fluorol yellow signal is detected in green and the xylem autofluorescence in blue (lignin); the corresponding brightfield micrographs (B, D, F, H, J, L) are also shown. Representative images from sections obtained from 8 up to 12 cm from the root tip with the typical observation with no suberized endodermis (A, G). Images showing that some specimens also presented a less frequent pattern, with few endodermal cells that also deposits suberin (C, I, E, K). In all observations, the exodermal cell layer was continuously suberized, except of individual unsuberized cells that corresponded to passage cells (marked with an asterisk). ex, exodermis; en, endodermis; xy, xylem. Scale bars correspond to 100 µm.

To determine the extent of suberin deficiency, the suberin monomeric composition of the roots grown in hydroponics was analyzed by gas chromatography (**Figure 3**). The amount of suberin was significantly affected by *CYP86A33* silencing. The total amount in the *CYP86A33*-RNAi roots decreased by 61.3% compared to that of wild type (wild type: 14.07 ± 0.74 μg mg^-1^; *CYP86A33*-RNAi: 5.45 ± 1.36 μg mg^-1^). This decrease in total suberin was due to a reduction of all different types of monomers, but especially ω-hydroxyacids and α,ω-diacids, which in *CYP86A33*-RNAi roots accounted for 40% and 22% of the wild type, respectively (**Figure 3A**). Although C16, C18, and C18:1 ω-hydroxyacids and their corresponding α,ω-diacids were the most reduced monomers in *CYP86A33*-RNAi compared to wild type, the other α,ω-functionalized monomers were also reduced (**Figure 3B**). Primary alcohols, fatty acids, and ferulic acids also decreased in *CYP86A33*-RNAi roots (**Figure 3C**).

**Figure 3.**
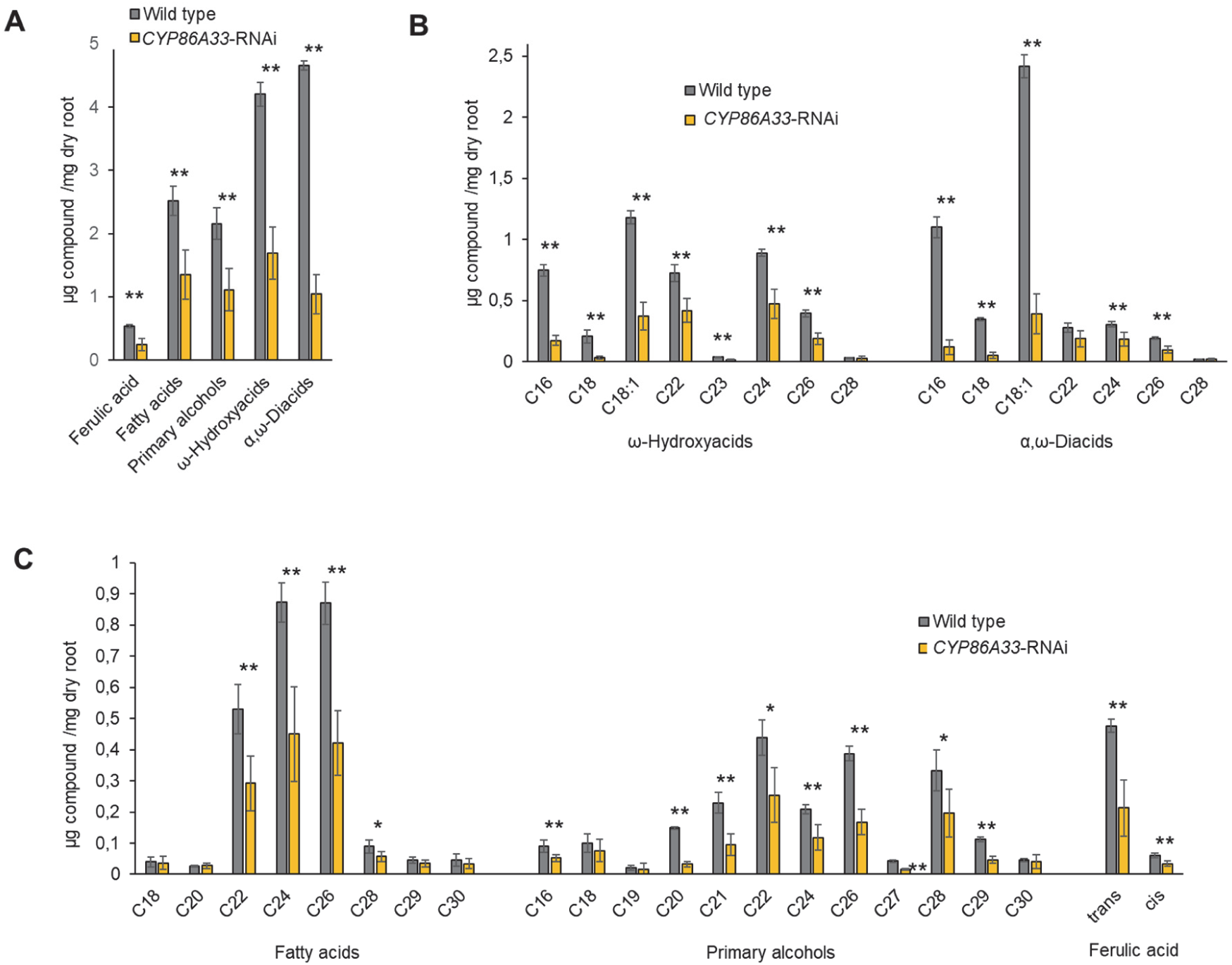
Chemical composition of root suberin preferentially deposited in exodermis in *CYP86A33*-RNAi silenced lines in comparison with wild type. The roots analyzed were from plants grown for three weeks in hydroponics. The relative amount of suberin monomers (µg / mg of dry root) grouped by compound class (A), the bi-functionalized monomers corresponding to ω-hydroxyacids and α,ω-diacids (B), and the fatty acids, primary alcohols and ferulic acids (C). Data are expressed as the mean ± SD of 3 wild type and 7 *CYP86A33*-RNAi biological replicates. The asterisks indicate statistically significant differences, t-test (* = p-value < 0.05: ** = p-value < 0.01).

### Effect of the exodermal suberin deficiency in plant nutrition

To study the relevance of exodermal suberin in root barriers to mineral nutrient transport, we compared the mineral element content of *CYP86A33*-RNAi plants, deficient in suberin, and those from wild-type (cv. Desirée) plants. *In vitro* plants were transferred to hydroponics and grown for three or seven weeks before analysis. The mineral nutrient content in the roots and shoots was quantified using inductively coupled plasma mass spectrometry (ICP). The results of element concentrations normalized to root or shoot dry weight were calculated (**Supplemental Tables 1-4**) and the data are summarized in a heatmap (**Figure 4**). Compared with the wild type, the ionome of *CYP86A33*-RNAi plants showed consistent changes at the two stages of plant development and in the two organs. Potassium (K) showed a significant decrease in roots and shoots of suberin-deficient mutant. All other significant differences observed in transgenic potatoes indicated a lower capacity of the suberin mutant to control the selective uptake of particular ions. Iron (Fe) higher uptake was observed at all stages and in both organs, although only significant at younger stages. Manganese (Mn), magnesium (Mg), copper (Cu), and sulfur (S) increased in all stages and organs of suberin-deficient mutants (S, data only seven-week plants), although the data were significant at specific stages. Other ions, such as sodium (Na) or zinc (Zn) revealed significant increases at specific plant stages, but the opposite trend (no statistically significant) was observed in the other plant stage. Overall, ionome analyses identified changes in metal content, indicating the importance of exodermal suberin in providing a selective barrier in the uptake or retention of mineral nutrients.

**Figure 4.**
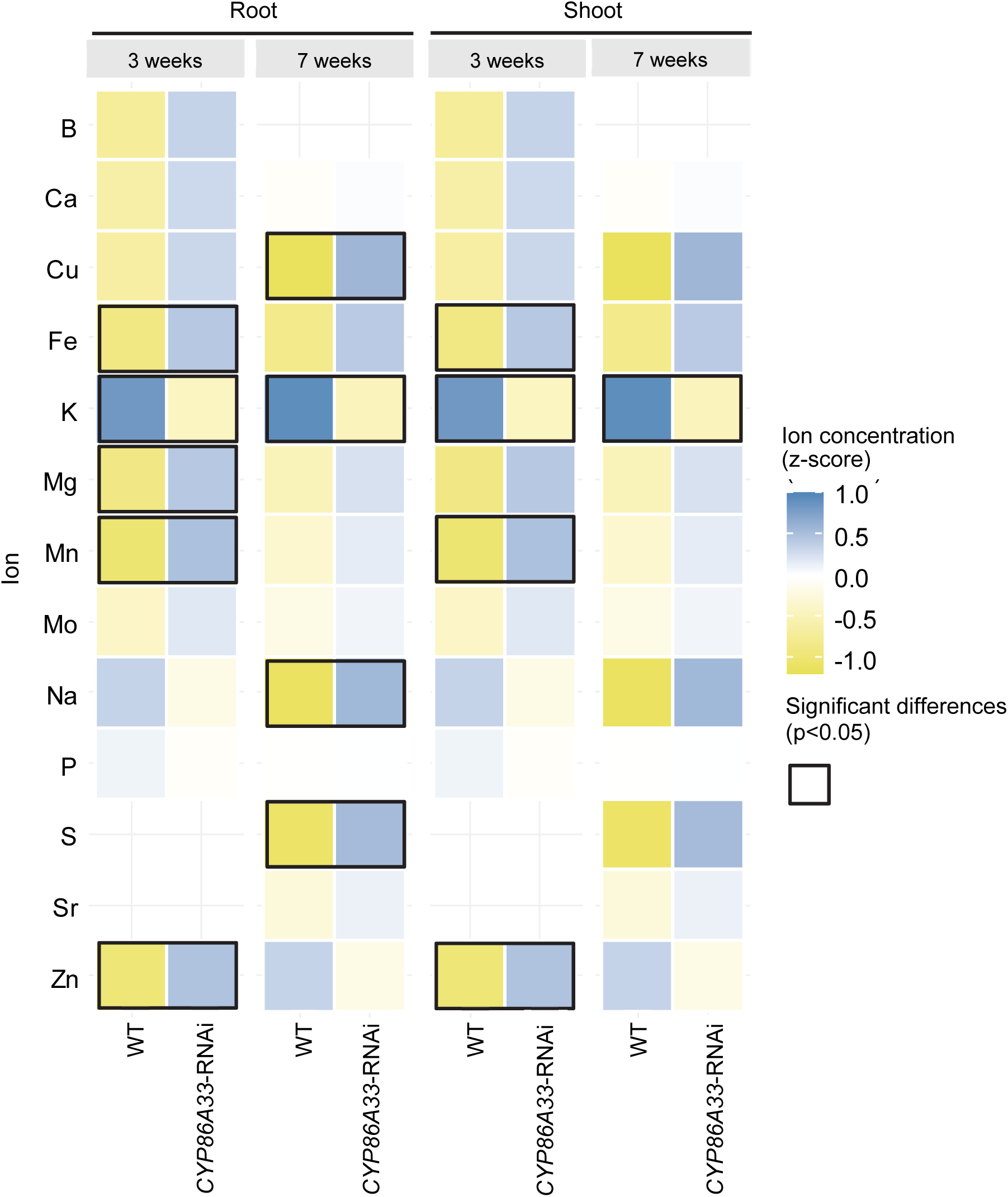
Effect of root suberin deficiency by *CYP86A33* downregulation in plant ionome. Heatmap showing the standardized mineral nutrient concentrations (μg / g sample dry weight) in root and shoot of *CYP86A33*-RNAi silenced and wild type (WT) plants grown in hydroponics for three and seven weeks. Significant differences (p < 0.05) in relation to suberin deficiency (*CYP86A33*-RNAi vs wild type) are outlined in black. Lines indicate elements that were not measured. The data corresponds to different biological replicates: 3-week grown roots: 4 WT and 8 *CYP86A33*-RNAi; 3-week grown shoots: 9 WT and 13 *CYP86A33*-RNAi; 7-week grown roots and shoots: 3 WT and 6 CYP86A33-RNAi, each.

### Spatial distribution of ions in potato roots

Since nutrient accumulation in root and shoot was dependent on the exodermal suberin present in adventitious roots, we next wanted to know: (i) the specific structural compartments that apoplastic barriers created in these potato roots, (ii) the compartment in which each mineral was retained, and (iii) whether exodermal suberin deficiency was able to change this nutrient distribution. To approach this, we mapped the spatial distribution of each individual element in the cross-sections of potato roots using micro Particle Induced x-ray Emission (micro-PIXE). Cryo-sections were obtained within 2 cm proximal to the root tip to include exodermal suberization and the absence of endodermal suberization (**Figure 1**). Micro-PIXE data was acquired for Na, Mg, P, S, K, Ca, Mn, Fe and Zn. In each mineral nutrient distribution map, atom localization showed clear differences across different anatomical compartments within the roots (**Figure 5**). In each elemental map, the regions comprising the recognized tissues were selected, and the concentration profile was extracted with GeoPIXE II software, as indicated by the rectangles in each of the micro-PIXE maps (**Figure 5**). To align these profiles for the element concentration across the tissue regions in the different cross-sections, micro-PIXE maps were manually adjusted to fit across the relative unit scale. Micro-PIXE data allows the mapping of atom localization across different tissue compartments (epidermis, cortex, central cylinder) generated by apoplastic barriers (exodermis and endodermis). The mappings showed four different groups of metal distribution: (i) K, and at less extent also P, had higher concentrations in the central cylinder; (i) Mg, Mn, and Zn had higher concentrations in the cortex (Mn also in the epidermis); (iii) Ca and S had similar distribution within the root; and (iv) Fe forms particular deposits in the epidermis/outer part of the root. This distribution was confirmed in two different replicates for each plant type, and the element concentrations of the tissues are presented in **Supplemental Table 5**. Despite being able to map the mineral nutrient distribution, wild type and *CYP86A33*-RNAi silenced line did not show significant changes across the root compartments. However, this is not rare considering that relatively small differences were observed in the bulk analyses of the whole root by ICP analyses and that higher variability is expected between root cross-sections due to technical limitations such as the difficulty in precisely cut at specific distances from the root tip (**Supplemental Table 1-4**).

**Figure 5.**
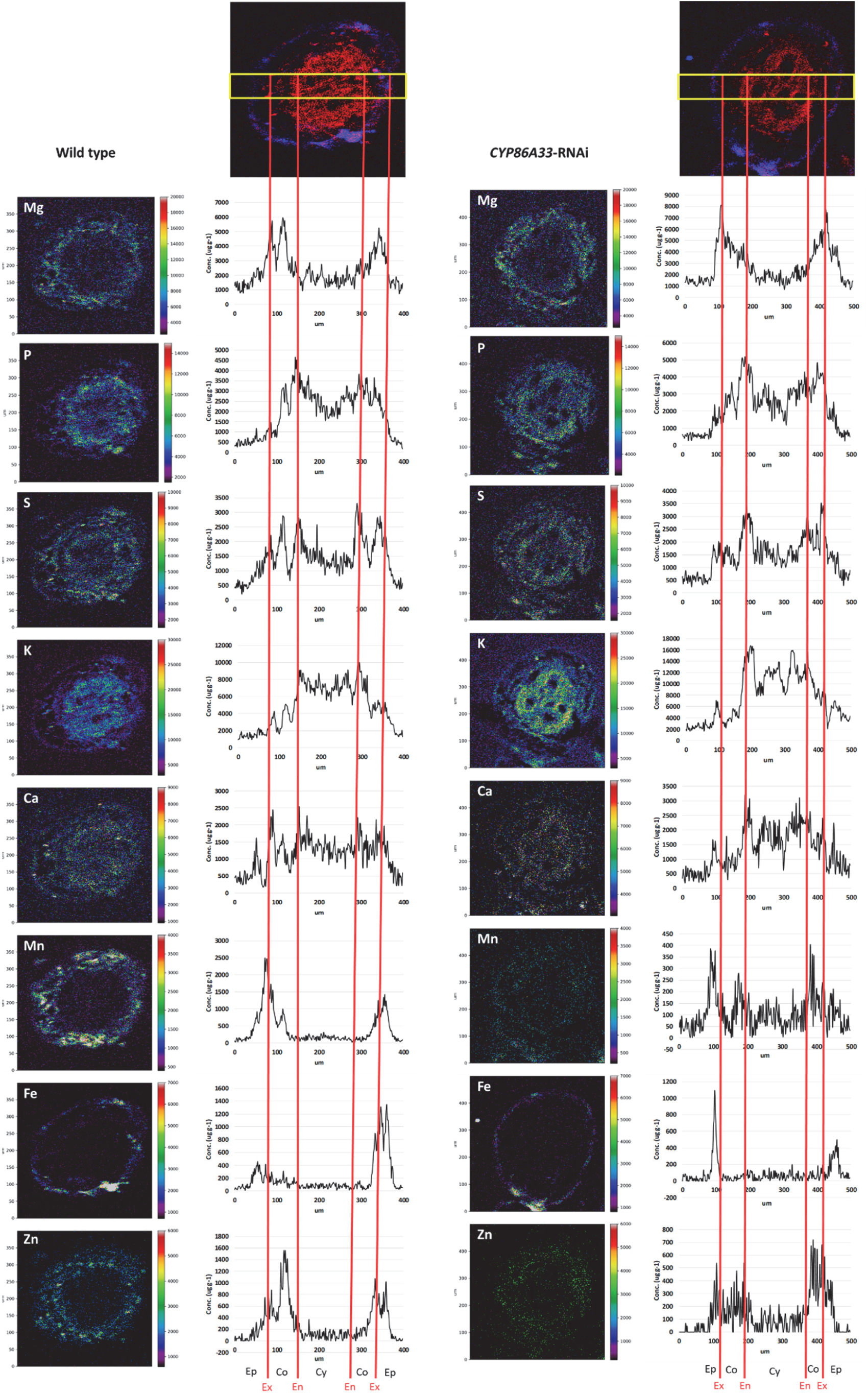
Quantitative element distribution maps of root cross-sections of wild-type and *CYP86A33*-RNAi potato roots grown in hydroponics. The yellow rectangle in the upper images of the section is 400 and 500 μm long, respectively, and defines the concentration profile analyzed for each nutrient element across the root. The red vertical lines indicate where the exodermis (Ex) and the endodermis (En) are located, defining the other structural compartments: epidermis (Ep), cortex (Co), central cylinder (Cy). For each element, the element distribution maps by micro-PIXE (in ppm) is shown on the left and the corresponding intensity in the rectangle defined in the upper images on the right.

### Silencing effects of *CYP86A33* in root and shoot biomass

Having identified that more than 61.3% (**Figure 3**) reduction of root suberization produced selective changes in the ionomic profile (**Figure 4**), we asked whether the suberin-induced changes in nutrition affected plant physiology. We first tested the effect on the growth of plants at two developmental stages: grown for three and seven weeks in hydroponics (**Figure 6**). Both shoots and roots of *CYP86A33*-RNAi plants accumulated significantly less biomass in shoots and roots (**Figure 6A**). Compared to the wild type, after three and seven weeks in hydroponics, the reduction in shoot biomass was roughly 21% and 16%, respectively, while root biomass was more severely affected and was reduced by 33% and 32%, respectively, as shown by the shoot/root biomass ratio (**Figure 6A**). We also investigated the importance of exodermal suberin in maintaining water within plants. Although the plants under analysis were grown in hydroponics for seven weeks, the water content in the aerial parts, leaves, and stems was significantly, but slightly, reduced in the suberin-deficient plants compared with the wild-type plants by 0.85% and 0.57%, respectively (**Figure 6B**). This slight defect in water retention in suberin-deficient mutant is unlikely to be relevant for plant growth but provides evidences on the functional role of exodermal suberin in restricting water movement. We did not observe changes in other physiological parameters such as the leaf water use efficiency, leaf transpiration, and leaf stomatal conductance (**Supplemental Figure 2**).

**Figure 6.**
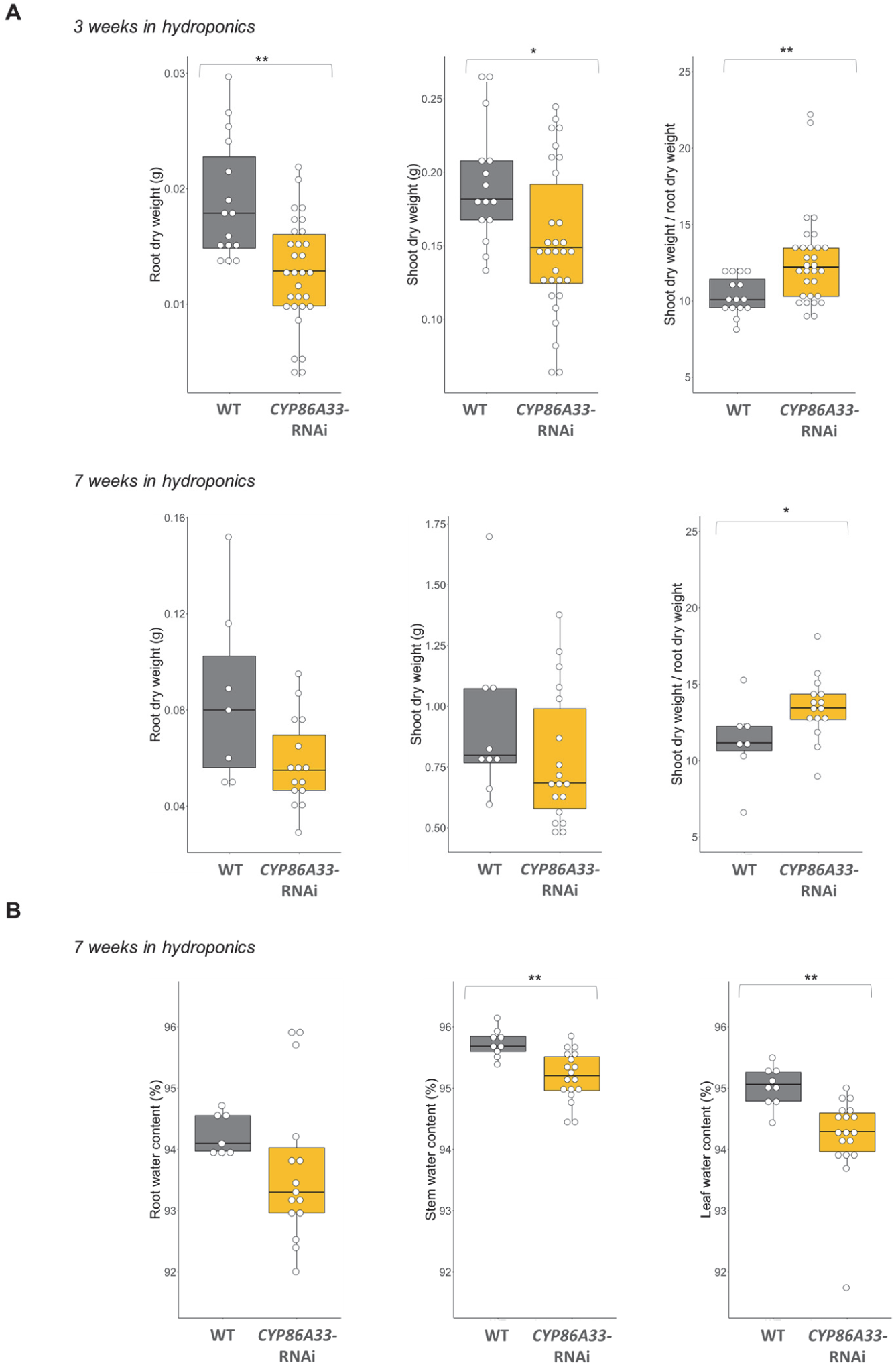
Effect of suberin-deficient CYP86A33-downregulated plants in plant biomass and water content. **A.** Boxplots of the root dry weight, shoot dry weight and shoot/root dry weights ratio of plants grown in hydroponics for three weeks (upper panel) and seven weeks (lower panel). **B.** Boxplots of the percentage of root, stem and leaf water content in plants grown for seven weeks in hydroponics. Individual replicates (n=7-30) are shown in the overlaid dotplots. The asterisks indicate statistically significant differences, t-test (* = p-value < 0.05; ** = p-value < 0.01).

## DISCUSSION

### Endodermis and exodermis of potato roots present a distinct spatiotemporal deposition of lignin and suberin

We have shown that potato root develops both endodermis and exodermis, single-layered tissues that modify their cell walls by accumulating lignin and suberin. The presence of the exodermis is not rare because most angiosperms (around 90%) develop this specialized hypodermis, which is defined to be very similar in structure to the endodermis, forms a Casparian strip, and quickly progresses to deposit suberin lamellae (Perumalla et al., 1990b; Enstone et al., 2003). Our observations indicate that endodermal and exodermal cells lignify in regions close to the root tip (below 2 cm); however, the lignification pattern differs between layers. While lignification in the endodermis forms the expected Casparian strip as a belt in the longitudinal direction, situated in the center of the anticlinal walls as typically observed (Meyer and Peterson, 2013; Geldner, 2013), lignification in the exodermis impregnates the anticlinal walls but extends to the outer tangential wall (**Figure 1A**). In fact, this lignification pattern was also identified in the closely related species *Solanum lycopersicum* (tomato) (Li et al., 2018) and *Glyceria maxima* (Soukup et al., 2007) and has recently been named lignified outer cap (Manzano et al., 2022). Even, Perumalla et al. (1990b) in their survey of hundreds of angiosperm species, commonly observed autofluorescence in the outer exodermal tangential wall (and epidermis), some persistent after hydrolyzing the suberin lamellae, suggesting that lignin impregnations in outer walls may be more common than previously expected.

Regarding suberization, we observed that exodermal cell progressed to state II of development by depositing the suberin lamellae, and formed a complete suberized cylinder surrounding the cortex close to the root tip (below 2 cm). This early ability of exodermis to fully suberize the complete cylinder is a common feature in potato adventitious roots as seen in different commercial varieties as well as a wild relative, regarding hydroponics or soil culture (**Figure 1B**, **Figure 1C**) (Łotocka et al., 2016). The early suberization of exodermis is also common for tomato or pepper (Cantó-Pastor et al., 2022), and this differs from other crop plants, suggesting a common pattern within Solanaceae family. In comparison, endodermal cell remains for longer (or even permanently) in the state I of differentiation and, in regions where exodermis suberization is fully completed, some endodermal cells, specifically those localized in the phloem pole, progress to state II of suberization (**Figure 1C, 1D, Figure 2**). Despite that the progression of endodermal suberization differs between cultivars, the advanced development of phloem-pole endodermis was generally observed, eventually leaving only unsuberized cells facing the xylem pole in some cultivars (passage cells) (**Figure 2**), as also observed previously (Łotocka et al., 2016). The early suberization process observed in the exodermis of potato adventitious roots calls into question the role of the endodermal suberization at root regions far from the root tip.

### The lignified endodermis and the ligno-suberized exodermis create anatomical compartments for nutrient accumulation and restrict their radial diffusion

The distribution of metal elements within the root tissues, namely the epidermis, cortex, and stele, at a distance of 2 cm from the root apex (**Figure 5**) suggests that anatomical barriers are established by the endodermis and exodermis through lignin and suberin deposition. The endodermis and exodermis in this region have already formed the Casparian strip and the polar outer lignification cap, respectively, creating two lignified longitudinal cylinder nets enclosing the cortex that effectively restrict the apoplastic movement of substances at the boundaries of the rhizodermis and stele (**Figure 1A**, **Figure 7**). Perumalla et al. (1990b), in their study of numerous angiosperm roots, found that roots developing an exodermis block apoplastic transport at exodermal radial walls, even when outer tangential lignification/suberization is extended to the outer exodermis domain or rhizodermis. In agreement, in tomato, a Solanaceae species similar to potato, it has been shown that the outer lignin cap in the exodermis is responsible for restricting the apoplastic movement across this layer (Manzano et al., 2022). Additionally, in this root region, the exodermis deposits suberin lamellae uniformly (**Figure 1**). This secondary cell wall layer of suberin is expected to impede apoplastic-to-influx transporter communication (or *vice versa*) at the plasma membrane, thus potentially blocking the transmembrane transport of nutrients, similar to the role of suberin in the endodermis (Barberon et al., 2016).

**Figure 7.**
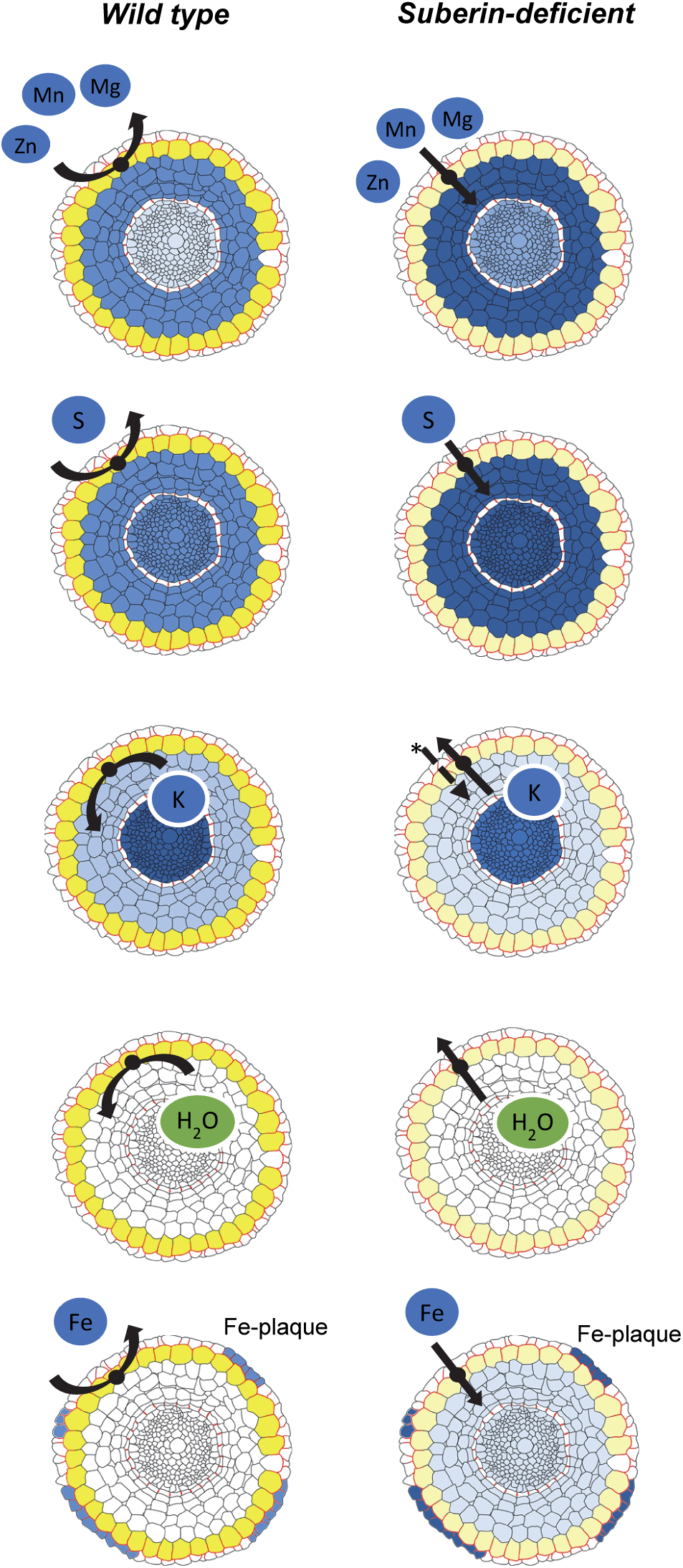
Schematic overview of the potato root integrating the data obtained related to the lignin and suberin deposits, the mineral nutrient distribution, and the effect of suberin deficiency on restricting radial transport across the exodermis. Each root cross-section integrates the histological detection of suberin and lignin: the high and low suberin amount in exodermis is painted in yellow and faint yellow, respectively, and lignin in endodermal Casparian strips and in exodermis radial and outer tangential walls is outlined in red. The mineral nutrient distribution by micro-PIXE analysis is shown in blue intensity and for each nutrient element the difference in blue intensity between the wild-type and the suberin-deficient panels indicates the concentration differences observed in the whole root by ICP analysis. The black arrows at the exodermis indicate the block of transport by suberin lamellae (wild type) or active transport at the exodermis due to suberin deficiency (suberin-deficient mutant). In the K suberin-deficient mutant panel, the asterisk highlights the possibility of lower K uptake by the suberin-deficient mutant.

By analyzing the mineral nutrient mappings in the root region characterized by the presence of the endodermal Casparian strip and the ligno-suberized exodermis, in conjunction with data from the *CYP86A33*-RNAi suberin-deficient mutant, several deductions concerning the radial movement of specific ions can be drawn (**Figure 7**). For instance, the increased levels of Mg, Mn, and Zn in the roots of the suberin-deficient mutant suggest that suberin defects facilitate the movement of these ions into and out of the exodermis more readily through influx and efflux transporters located at the exodermis plasma membrane. Specific transporters for these ions have been identified for the exodermis in rice, where OsNramp5 and OsMTP9 are polarly localized at the outer and inner side of the exodermis, respectively, and play a major role in Mn uptake (Sasaki et al., 2016). The accumulation of Mn, Mg, and Zn ions in the root cortex (**Figure 5**, **Figure 7**) further indicates that the Casparian strip network in the endodermis primarily impedes their radial movement to the stele. However, the higher increase in Zn and Fe in the shoots of the *CYP86A33*-RNAi mutant suggests a more efficient transport through the endodermis, which would potentially maintain the transcellular or symplastic pathway active. Notably, in Arabidopsis, the influx transporters AtYSL2 and AtIRT3, which are localized in endodermis, are known to contribute to the uptake of both Fe and Zn (Schaaf et al., 2005; Lin et al., 2009; Bao et al., 2019). Further investigation is required to understand whether regulatory or compensatory mechanisms at the endodermis contribute to this higher translocation to the shoot.

The iron aggregates observed in the outer part of the root (**Figure 5**) probably correspond to iron plaque deposits. Iron plaques are Fe^3+^ oxyhydroxide precipitates commonly observed in plants grown in hydroponics or under waterlogged conditions, such as rice, resulting from Fe^2+^ (ferrous iron) oxidation by radial oxygen loss (ROL) leaking from the root (Maisch et al., 2020). Suberin has been described as a barrier to ROL (Ejiri and Shiono, 2019), so the higher accumulation of Fe ions in the suberin-deficient potato roots could involve an increased oxygen leakage through the suberin-deficient exodermis, promoting iron plaque formation in hydroponics, similar to what is observed in rice (Wu et al., 2012) (**Figure 7**). Additionally, the higher accumulation of iron in the shoots of the suberin mutant indicates a higher uptake of iron, which in dicotyledonous involves ferric chelate reductases (FRO2) on the plasma membrane, reducing Fe^3+^ to the more soluble Fe^2+^, subsequently absorbed by the major iron transporter, IRT1 (Eide et al., 1996; Robinson et al., 1999).

As for K and P, their retention in the central cylinder indicates that these ions cannot backflow through the unsuberized endodermis apoplastically due to the Casparian strip network (**Figure 5**, **Figure 7**). However, the substantial decrease in K levels in the roots and shoots of the suberin-deficient mutant implies that either a) lower K uptake or b) higher backflow of K through the endodermis and eventually exodermis is needed to reach the rhizosphere. In the latter scenario, efflux transporters or symplastic backflow at the endodermis and exodermis would be expected. Nonetheless, compensatory effects for K are also plausible, considering that K^+^ plays a crucial role in maintaining ion homeostasis and physiological stability (Srivastava et al., 2020; Mostofa et al., 2022).

### Both the endodermal and exodermal suberin contribute to maintaining nutrient homeostasis, playing a pivotal role in plant growth

The role of suberin in controlling ion homeostasis has been previously described for endodermis in suberin-deficient mutants of Arabidopsis (Barberon et al., 2016; Wang et al., 2020). In this endodermis model, suberin deficiency leads to increased uptake of calcium, manganese, and sodium, while sulfur and potassium uptake are reduced. This is in part in agreement with our data where the exodermal suberin is much affected (**Figure 4**). In our suberin deficient exodermal model, the uptake of manganese, zinc, copper, and magnesium is enhanced, and potassium is severely down-accumulated in the entire plant. These similarities demonstrated that exodermis can function similarly to endodermis and contribute to nutrient homeostasis and suggest that the uptake of these nutrients likely follows a coupled transcellular pathway involving influx and efflux transporters located at each tangential domain of the plasma membrane of the exodermal cells, paralleling those at endodermal cells (Barberon and Geldner, 2014; Bao et al., 2019). Additionally, we also observed that iron uptake was higher in suberin-deficient potato roots indicating that suberin blocks its uptake at the exodermal layer. The suberin ability to block iron uptake agrees with the Fe deficiency capacity to delay the endodermal suberization as does the *irt1* iron uptake transporter loss-of-function mutant (Barberon et al., 2016).

Although the alteration of nutrient homeostasis in the suberin-deficient mutant is not severe, especially in shoots, the negative consequences on plant growth are significant, with a more pronounced effect on root growth (**Figure 6**), likely due to more pronounced alterations in the ionome of roots (**Figure 4**). This is not surprising considering that various minerals participate in essential and diverse biochemical processes, serving pivotal functions in plant biology (Mulet et al., 2020). As an example, K^+^, which is the most affected nutrient in the suberin-deficient mutant, regulates numerous physiological processes directly impacting plant growth (Mostofa et al., 2022). Overall, the data underscore the importance of exodermal suberin’s ability to regulate nutrient homeostasis in plant growth, as also demonstrated in Arabidopsis endodermis (Barberon et al., 2016). Remarkably, even under hydroponic conditions, the impact of suberin on water retention is evident, as demonstrated by the suberin-deficient mutant exhibiting a slight reduction in leaf water content (**Figure 6**).

## METHODS

### Plant material and growth conditions

Potato plants (*Solanum tuberosum* L.) subsp*. tuberosum* cv Desirée and cv. Red Pontiac, and a wild relative *Solanum tuberosum* subsp. *andigena* were used to identify the suberization pattern of the root apoplastic barriers, exodermis and endodermis. In cv. Desirée background, the potato plant *CYP86A33-RNAi* (line 22 and 39) has downregulated by RNAi the c*ytochrome P450 fatty acid ω-hydroxylase CYP86A33*, which is orthologous of the Arabidopsis *CYP86A1/HORST* (At5g58860) (Bjelica et al., 2016), and has a 60% reduction of suberin in potato tuber phellem (Serra et al., 2009).

For the growth in hydroponics, we used plants grown in vitro in a solid MS (Murashige and Skoog) media supplemented with 2% sucrose (2MS) for 2 up to 4 weeks. These plants were transferred to aerated half-strength Hoagland solution in a 10-L volume buckets for three weeks (younger plants) and seven weeks (older plants) before subsequent analysis. The nutrient solution was changed every seven days. The nutrient solution was based on Hoagland growth medium with the following concentrations: 2.5 mM KNO_3_, 2.5 mM Ca(NO_3_)_2_, 12 µM 6% Fe EDDHA (Kelamix Fe, Burés Professional), 1 mM MgSO_4_ꞏ7H_2_O, 0.5 mM NH_4_NO_3_, 23 µM H_3_BO_3_, 4.2 µM MnCl_2_ꞏ4H_2_O, 3.8 µM ZnSO_4_ꞏ7H_2_O, 0.14 µM CuSO_4_ꞏ5H_2_O, 0.25 µM Na_2_MoO_4_ꞏ2H_2_O and 0.25 mM KH_2_PO_4_. Plants *in vitro* and plants in hydroponics were grown in chambers under a light/dark photoperiod cycle of 12 h/12 h at 24 °C and 22 °C, respectively. For the histochemical root analyses, we also analyzed the roots of potatoes grown in soil at the same photoperiodic and temperature conditions.

### Histochemical analysis of suberin staining

For fluorol yellow staining, roots from plants grown in soil and hydroponics were cut in 2-cm segments and stored in methanol before staining. Root segments were first counter-stained with aniline blue (5% w/v, in H_2_O) (Fluka Chemie) for 30 s and thereafter were washed in distilled water to remove the excess stain. This allowed to reduce the background autofluorescence and to help to visualize the root during free-hand sectioning. Aniline blue-stained roots were then embedded into fresh 6% aqueous melt agarose (Agarose D1 low EEO, CONDALAB) kept at 53 °C in a thermoblock. Free-hand root cross-sections obtained with sharp razor blades were placed into distilled water and subsequently stained with fresh fluorol yellow 088 (0.01% w/v, in methanol) (Sigma-Aldrich, Merck) at room temperature for 1 h in darkness and rinsed and kept into distilled water before observation. Sections were mounted on glass slides in distilled water and observed on an Olympus Vanox-T AH2 epifluorescence microscope using a UV excitation filter (excitation at 330-389 nm) (otherwise stated), collecting emission fluorescence from 420 nm. Micrographs were acquired using an Olympus DT73 digital camera and the Olympus CellSens Standard Software (v.1.11), and were finally processed using Fiji-ImageJ software (v.1.23).

For Nile red staining, the 2-cm root segments grown in hydroponics were fixed, cleared and stained as previously described by Ursache et al. (2018) with modifications. Briefly, roots were fixed in 4% paraformaldehid in 1x PBS buffer for 60 min under vacuum and rinsed twice for 1 min in 1x PBS buffer. Root samples were then cleared with ClearSee (10% xylitol, 15% sodium deoxycholate, 25% urea in distilled water) for at least 3 days, and fresh Clearsee solution was substituted every week. For Nile red staining, we first stained the root segments with 0.1% of calcofluor white (Sigma-Aldrich) in ClearSee for two days for general polysaccharide staining, then samples were rinsed with ClearSee for 30 min, stained with 0.05% of Nile red in ClearSee for three days and rinsed in ClearSee for at least 30 min before observation under an inverted NIKON A1R confocal microscope. All the incubations were done at room temperature, darkness and in agitation. The excitation and emission spectra for calcofluor white were 405 nm and 425-475 nm, respectively; for Nile red 543.5 nm and 570-620 nm, respectively. Images were obtained with NIS-Elements Viewer software (Nikon) and processed using Fiji-ImageJ software (v.1.23).

### RNA extraction, cDNA synthesis and Real time RT-PCR analyses

Total RNA was isolated from root tissue of 4 biological replicates (individual plants) following the guanidine hydrochloride method (Logemann et al., 1987). RNA was treated with DNA-free DNase Treatment and Removal Reagents (Ambion, Life Technologies) and cDNA was syntethized from 1 µg of RNA using the High Capacity cDNA Reverse Transcription kit (Applied Biosystems). For the real-time RT-PCR analysis, forward and reverse primer sequences were, respectively: 5′-TCTACTGGGGTATCCGCAAC-3′ and 5′-TTTGGTGAAAGGGTTTCAGG-3′ for *CYP86A33* gene, and 5’-GAACCGGAGCAGGTGAAGAA-3’ and 5’-GAAGCAATCCCAGCGATACG-3’ for the reference gene *adenine phosphoribosyl transferase* (*APRT*). Real-time PCR analysis was performed using a LightCycler® 96 Real-Time PCR System (Roche). RT-qPCR reaction was prepared in triplicates by mixing 2.5 μl of a 25-fold diluted cDNA, 300 nM each of forward and reverse primers, 5 μl of SYBR Green Select Master Mix (Roche), and, and up to 10 µl with RNAse free water. The thermocycler conditions were 95 °C for 10 min; 40 cycles of 95 °C for 10 s and 60 °C for 60 s, followed by a final dissociation step to confirm a single amplicon. The efficiency (E) for each primer pair was calculated using five dilutions of template and the equation E=10^(−1/slope)^. Relative transcript accumulation (RTA) was calculated as = (E_target_)^(Ct_control - Ct_sample)^ / (E_reference_)^Ct_control - Ct_sample)^ (Pfaffl, 2001), being the control one of the wild-type samples and the reference the housekeeping gene ACT. Three Triplicates of four biological replicates (roots of individual plants) (n=4) were analyzed for each genotype.

### Suberin chemical composition of plant roots

For the isolation of suberized barriers, each individual root biological replicate included the roots from 3 plants. Three wild-types and 7 *CYP86A33*-RNAi (n=3 line 22, n=4 line 39) biological replicates were used. Plant roots were treated as described previously (Company-Arumí et al., 2016). In detail, after harvesting, roots were washed with distilled water and treated at room temperature for four weeks with a mixture of 5% v/v cellulase and 5% v/v pectinase diluted in citric buffer (10^-2^ M pH 3.0, adjusted with KOH) and 1 mM sodium azide to prevent bacterial growth. Then, tissues were treated for one day each with boric acid buffer (2 x 10^-2^ M pH 9.0) and deionized water, and they were dried and stored in the dark until use. The isolated material from one root (1-2 mg) was treated using 2 mL of chloroform:methanol mixture (1:1 v/v) over a period of 16-18 hours at 50 rpm and then rinsed three times with the mixture to remove the remaining wax material.

Suberin was depolymerized by transesterification by immersing the wax-free residues in boron trifluoride in methanol (10% BF_3_/MeOH) and incubating the samples at 70 °C for 16-18 hours in a Teflon-sealed screw-cap tube. After the reaction took place, 10 µg of the dotriacontane were added as a surrogate and the methanolysate was transferred to a new vial containing 2 mL of saturated NaHCO_3_ aqueous solution. The solid residue was rinsed twice with chloroform and the cleaning solutions were added to the methanolysate and NaHCO_3_ mixture. The aqueous-methanol phase was then extracted twice with chloroform and, after phase separation, the chloroform extract was rinsed with ultrapure water. Anhydrous sodium sulphate powder was added to the organic phase to remove traces of water and the solvent was then evaporated to dryness. The released monomers were transformed to tms derivatives by BSTFA derivatization, adding 20 μL of the reagent and 20 μL of pyridine to the dry residues and incubating the samples for 40 min at 70 °C. An appropriate volume of chloroform was added to the final solutions to obtain the desired concentrations for gas chromatography (GC) analyses.

GC-mass spectrometry (GC-MS) was used for suberin monomer tms derivative identification and GC-FID for its quantification. GC-flame ionization detection (GC-FID) analysis was performed in a Shimadzu GC-2010 Plus using a BP1 capillary column (30 m length, 0.25 mm i.d., 0.1 µm film thickness, Teknokroma). A split/ splitless injector was used in the splitless injection mode (splitless time 1 min) with the injector temperature at 280 °C. Helium was used as the carrier gas at a constant flow rate of 1 mL/min. The detector temperature was maintained at 320 °C. The initial oven temperature program started at 140 °C, followed by 3 °C min^-1^ increases up to 310 °C, where it was held for 5 min. Chromatograms were processed using GC Solution software (version 2.41) from Shimadzu. GC-MS analysis, performed using a selective mass detector with ion trap (Trace GC 2000 series coupled to a Thermo Scientific Polaris Q mass spectrometer), enabled the identification of the derivatized suberin monomers comparing the mass spectra with available standards or literature (Kolattukudy and Agrawal, 1974; Zeier and Schreiber, 1997, 1998).

### ICP analyses for mineral nutrient chemical composition

Roots and shoots collected from hydroponics were dried in an oven at 70 °C for 72 h and thereafter were weighted on an analytical balance to obtain the dry weight. Each individual root biological replicate included pools of roots from 1 to 3 plants grown in the same bucket and each shoot replicate included pools of shoots from 1 to 2 plants, to reach the minimum tissue mass for ICP analysis. Dried samples (0.08 - 0.12 g) were cut into small pieces and placed in PTFE digestion vessels. Acid digestion was performed by adding 9 mL of 69% nitric acid (HNO_3_) (Suprapur, Merck) and 1 mL of 30% hydrogen peroxide (H_2_O_2_) (Trace Select, Fluka). Vessels were capped and heated into the microwave digestion system (Speedwave XPERT, BERGHOF, Germany) following the program: 5 min to reach 180 °C and 10 min at 180 °C. After cooling, digested sample solutions were transferred to 25 mL vials and brought to volume with ultrapure de-ionized water. Samples were stored at 4 °C prior to analysis.

Nutrient element content (ionome studies) was measured using inductively coupled plasma atomic emission spectrometry (ICP-OES) or ICP-mass spectrometry (ICP-MS). For ICP-OES an Agilent Technology model Vertical Dual View 5100 ICP-OES spectrometer equipped with an SPS3 autosampler. The instrument was fitted with a SeaSpray® concentric glass nebulizer, a double-pass cyclonic spray chamber and an easy-fit one-piece torch with a 1.8 mm id injector. The detector type was a CCD (charge-coupled device). For ICP-MS, a quadrupole-based ICP-MS system (Agilent 7500c, Agilent Technologies) was used, equipped with an octopole collision reaction cell. ^103^Rh was used as internal standard. The accuracy of the analysis was checked by concurrent analysis of standard reference materials. Equipment calibration was performed using multi-element calibration standards prepared from single-element standard solutions (1000 mg L^-1^) (Pure Chemistry). Sample concentrations were calculated using an external calibration method and the values were normalized to the amount of the sample processed. Each sample was run in triplicate and the mean was considered the representative value for a particular sample. For the root analysis of younger plants (three weeks in hydroponics), 4 wild-type and 8 *CYP86A33*-RNAi (n=4 line 22, n=4 line 39) biological replicates were used; for the shoot analysis 9 wild-type and 13 *CYP86A33*-RNAi (n=6 line 22 and n=7 line 39). For both the root and shoot analyses of older plants (seven weeks in hydroponics) 3 wild-types and 6 *CYP86A33*-RNAi (n=3 line 22, n=3 line 39) biological replicates were used. Trends of the ionomic profiles were presented in a heatmap, displaying the concentration per each mineral nutrient after applying z-scores transformations, in roots and shoots respectively, using the ggplot2 package (v.3.3.4; https://ggplot2.tidyverse.org/) in R.

### Mapping of element distribution by micro-proton induced X-ray emission analysis (micro-PIXE)

The 2 cm of the root tip was cut and introduced into an stainless steel needle with an inner diameter of 2 mm, leaving around 2 mm of the root outside the needle which was submerged in a tissue-freezing medium (Jung, Leica) and quickly frozen in liquid nitrogen (Vogel-Mikuš et al., 2014). Cryo-sections of 50 µm thickness were obtained at −25 °C using a Leica CM3050 cryotome. Sections were placed in custom-made aluminium holders and covered with a stainless-steel fitting cover to keep the section flat. Using a cryo-transfer assembly cooled by liquid nitrogen, the sections were freeze-dried for 3 days at −25 °C and at 0.240 mbar in a freeze dryer (Alpha 2-4 Christ, Osterode am Harz, Germany). Dry sections were placed between two thin layers of Pioloform foil stretched on aluminium holders and imaged under bright field and UV excitation (365 nm) using Zeiss Axioskop 2 MOT microscope equipped with an Axiocam MRc colour digital camera (Vogel-Mikuš et al., 2014; Pongrac et al., 2019).

Root cross-sections were used to measure the tissue-specific distributions of the different elements by micro-particle-induced X-ray emission (micro-PIXE). Micro-PIXE analysis was performed at the nuclear microprobe of the Jožef Stefan Institute as previously described (Detterbeck et al., 2016; Lyubenova et al., 2013; Vavpetič et al., 2015). To determine beam exit energy from the sample, related to the sample local tissue density, an on-off axis scanning transmission ion microscopy (STIM) was simultaneously performed (Pallon et al., 2004; Vavpetič et al., 2013). From micro-PIXE spectra we calculated numerical matrices (pixel-by-pixel concentration matrices) and generated distribution and co-localization maps using GeoPIXE II software package (Ryan, 2000) utilizing the dynamic analysis method (Ryan et al., 2015). To further enhance image contrast we applied smooth (Gaussian function, standard deviation 1.5) and edge enhance (Roberts function) filters. Using ImageJ, the areas enclosed between tissues (epidermis, cortex and central cylinder) were defined and the concentrations for each element were extracted (Singh et al., 2014; Vogel-Mikuš et al., 2014).

### Physiological parameters: biomass and leaf water content

Wild-type and suberin-deficient mutant (line 39 and line 22) plants were grown in hydroponics. For each plant (biological replicate), leaves, stems and roots were separated, being roots washed with distilled water and the remaining surface watered quickly and lightly dried. Then, plant fractions were weighed to determine their fresh mass (FM). To determine the dry mass (DM), plant fractions were oven-dried at 60 °C for 3 days and weighted. The water content of leaves (LWC), stems (SWC) and roots (RWC) were calculated as: WC (%) = (FM-DM) x 100/FM.

Foliar gas exchange parameters, including transpiration rates (E), stomatal conductance (gs) and photosynthesis were measured in one attached leaf for plant (biological replicate) using a portable open-circuit infrared gas analyzer system (CIRAS-2, PP-Systems Inc. Amesbury, USA). Intrinsic water use efficiency was calculated as A/gs (WUE).

### Statistical analyses

The data were compared based on the mean values using a t-test for independent samples and significance was considered when p < 0.05. Data that do not meet variance homogeneity by Levene’s test (p < 0.05) was analyzed using the non-parametric Mann-Whitney test (p < 0.05) and when significant it was indicated specifically. Normal distribution was assumed.

### Accession Numbers

*CYP86A33* potato gene from this article can be found in the EMBL/GenBank data libraries under accession number EU293405.

## Supplemental Data files

**Supplemental Figure 1.** C*Y*P86A33 transcript accumulation in the roots of wild-type and RNAi-silenced potato plants.

**Supplemental Figure 2.** Effect of suberin-deficient *CYP86A33-*downregulated plants in physiological parameter performance.

**Supplemental Table 1.** Nutrient element concentration in the root of *CYP86A33*-silenced and wild-type plants grown for three weeks in hydroponics.

**Supplemental Table 2.** Nutrient element concentration in the shoot of *CYP86A33*-silenced and wild-type plants grown for three weeks in hydroponics.

**Supplemental Table 3.** Nutrient element concentration in the root of *CYP86A33*-silenced and wild-type plants grown for seven weeks in hydroponics.

**Supplemental Table 4.** Nutrient element concentration in the shoot of *CYP86A33*-silenced and wild-type plants grown for seven weeks in hydroponics.

**Supplemental Table 5.** Element concentrations in dpidermis, cortex and central cylinder (cylinder) of the potato roots grown in hydroponics for three weeks, obtained by micro-PIXE and analyzed using the GeoPIXE II software.

## ACKNOWLEDGEMENTS

We are grateful to Iván Herrero for staining the potato root material. This work was supported by the Ministerio de Economía y Competitividad [AGL2012-36725; AGL2015-67495-C2-1-R (MINECO/FEDER,UE)] and Ministerio de Ciencia e Innovación [PID2019-110330GB-C21(MCI/ AEI)] and The European Regional Development Fund (ERDF). D. Company-Arumí was supported by a University of Girona PhD fellowship (BR10/27) and C. Montells by a Spanish Research Collaboration fellowship (Ministerio de Educación y Formación Profesional). We acknowledge a financial support from ARIS, P1-0212 Plant biology program group.

## AUTHOR CONTRIBUTIONS

E.A. and O.S. conceived and designed the research. M.F, O.S. and E.A. obtained the financial support. Different authors performed the analyses and provided the data: D.C.-A., C.M., M.I, E.A. and E.M. ionomic analyses, D.V. and C.M. physiological analyses, D.C.-A, E.A. and O.S. suberin chemical analyses, K.V.-M. and M.K. PIXE-analyses, C.M. and O.S. histological analyses. Data analyses and interpretation was performed by all authors. O.S. wrote the manuscript with the input of E.A.; O.S., C.M. and K.V.-M. made the figures, and all the authors revised the final manuscript form.

## REFERENCES

Bao, Z., Bai, J., Cui, H., and Gong, C. (2019). A Missing link in radial ion transport: Ion transporters in the endodermis. Front Plant Sci 10: 713.

Barberon, M. and Geldner, N. (2014). Radial transport of nutrients: the plant root as a polarized epithelium. Plant Physiol 166: 528–37.

Barberon, M., Vermeer, J.E.M., De Bellis, D., Wang, P., Naseer, S., Andersen, T.G., Humbel, B.M., Nawrath, C., Takano, J., Salt, D.E., and Geldner, N. (2016). Adaptation of root function by nutrient-induced plasticity of endodermal differentiation. Cell 164: 1–13.

Bjelica, A., Haggitt, M.L., Woolfson, K.N., Lee, D.P.N., Makhzoum, A.B., and Bernards, M.A. (2016). Fatty acid ω-hydroxylases from *Solanum tuberosum*. Plant Cell Rep 35: 2435–2448.

Calvo-Polanco, M. et al. (2021). Physiological roles of Casparian strips and suberin in the transport of water and solutes. New Phytol 232: 2295–2307.

Cantó-Pastor, A. et al. (2022). A suberized exodermis is required for tomato drought tolerance. bioRxiv 2022.10.10.511665; doi: 10.1101/2022.10.10.511665: 2022.10.10.511665.

Company-Arumí, D., Figueras, M., Salvadó, V., Molinas, M., Serra, O., and Anticó, E. (2016). The Identification and quantification of suberin monomers of root and tuber periderm from potato (*Solanum tuberosum*) as aatty acyl *tert* -butyldimethylsilyl derivatives. Phytochem Anal 27: 326–335.

Damus, M., Peterson, R.L., Enstone, D.E., and Peterson, C.A. (1997). Modifications of cortical cell walls in roots of seedless vascular plants. Botanica Acta 110: 190–195.

Detterbeck, A., Pongrac, P., Rensch, S., Reuscher, S., Pečovnik, M., Vavpetič, P., Pelicon, P., Holzheu, S., Krämer, U., and Clemens, S. (2016). Spatially resolved analysis of variation in barley (*Hordeum vulgare*) grain micronutrient accumulation. New Phytol 211: 1241–1254.

Eide, D., Broderius, M., Fett, J., and Guerinot, M.L. (1996). A novel iron-regulated metal transporter from plants identified by functional expression in yeast. Proc Natl Acad Sci U S A 93: 5624–5628.

Ejiri, M. and Shiono, K. (2019). Prevention of radial oxygen loss is associated with exodermal suberin along adventitious roots of annual wild species of Echinochloa. Front Plant Sci 10: 254.

Enstone, D.E., Peterson, C. A., and Ma, F. (2003). Root endodermis and exodermis: structure, function, and responses to the environment. J Plant Growth Reg 21: 335–351.

Geldner, N. (2013). The endodermis. Annu Rev Plant Biol 64: 531–558.

Hose, E., Clarkson, D.T., Steudle, E., Schreiber, L., and Hartung, W. (2001). The exodermis: a variable apoplastic barrier. J Exp Bot 52: 2245–2264.

Joshi, M. and Ginzberg, I. (2021). Adventitious root formation in crops—Potato as an example. Physiologia Plantarum 172: 124–133.

Kajala, K. et al. (2021). Innovation, conservation, and repurposing of gene function in root cell type development. Cell 184: 3333–3348.e19.

Kolattukudy, P.E. and Agrawal, V.P. (1974). Structure and composition of aliphatic constituents of potato tuber skin (suberin). Lipids 9: 682–691.

Kreszies, T., Schreiber, L., and Ranathunge, K. (2018). Suberized transport barriers in Arabidopsis, barley and rice roots: From the model plant to crop species. J Plant Physiol 227: 75–83.

Li, B., Kamiya, T., Kalmbach, L., Yamagami, M., Yamaguchi, K., Shigenobu, S., Sawa, S., Danku, J.M.C., Salt, D.E., Geldner, N., and Fujiwara, T. (2017). Role of LOTR1 in nutrient transport through organization of spatial distribution of root endodermal barriers. Curr Biol 27: 758–765.

Li, P., Yang, M., Chang, J., Wu, J., Zhong, F., Rahman, A., Qin, H., and Wu, S. (2018). Spatial expression and functional analysis of Casparian strip regulatory genes in endodermis reveals the conserved mechanism in tomato. Front Plant Sci 9: 832.

Lin, Y.-F., Liang, H.-M., Yang, S.-Y., Boch, A., Clemens, S., Chen, C.-C., Wu, J.-F., Huang, J.-L., and Yeh, K.-C. (2009). Arabidopsis IRT3 is a zinc-regulated and plasma membrane localized zinc/iron transporter. New Phytol 182: 392–404.

Líška, D., Martinka, M., Kohanová, J., and Lux, A. (2016). Asymmetrical development of root endodermis and exodermis in reaction to abiotic stresses. Ann Bot 118: 667–674.

Łotocka, B., Kozak, M., and Rykaczewska, K. (2016). Morphology and anatomy of the root system of new potato cultivars. Part II. Root anatomy. Biuletyn IHAR: 31–43.

Lyubenova, L., Pongrac, P., Vogel-Mikuš, K., Mezek, G.K., Vavpetič, P., Grlj, N., Regvar, M., Pelicon, P., and Schröder, P. (2013). The fate of arsenic, cadmium and lead in Typha latifolia: A case study on the applicability of micro-PIXE in plant ionomics. J Hazard Mat 248–249: 371–378.

Maisch, M., Lueder, U., Kappler, A., and Schmidt, C. (2020). From plant to paddy—how rice root iron plaque can affect the paddy field iron cycling. Soil Syst 4: 28.

Manzano, C. et al. (2022). Regulation and Function of a Polarly Localized Lignin Barrier in the Exodermis. bioRxiv 2022.10.20.513117; doi: 10.1101/2022.10.20.513117

Meyer, C.J. and Peterson, C.A. (2013) Structure and Function of Three Suberized Cell Layers: Epidermis, Exodermis, and Endodermis. In Plant Roots, The Hidden Half CRC Press.

Mostofa, M.G., Rahman, Md.M., Ghosh, T.K., Kabir, A.H., Abdelrahman, M., Rahman Khan, Md.A., Mochida, K., and Tran, L.-S.P. (2022). Potassium in plant physiological adaptation to abiotic stresses. Plant Physiol Biochem 186: 279–289.

Mulet, J.M., Campos, F., and Yenush, L. (2020). Editorial: Ion Homeostasis in Plant Stress and Development. Front Plant Sci 11: 618273.

Namyslov, J., Bauriedlová, Z., Janoušková, J., Soukup, A., and Tylová, E. (2020). Exodermis and endodermis respond to nutrient deficiency in nutrient-specific and localized manner. Plants (Basel) 9: E201.

Pallon, J., Auzelyte, V., Elfman, M., Garmer, M., Kristiansson, P., Malmqvist, K., Nilsson, C., Shariff, A., and Wegdén, M. (2004). An off-axis STIM procedure for precise mass determination and imaging. Nuclear Instruments and Methods in Physics Research Section B: Beam Interactions with Materials and Atoms 219–220: 988–993.

Perumalla, C.J., Chmielewski, J.G., and Peterson, C.A. (1990a). A survey of angiosperm species to detect hypodermal Casparian bands. III. Rhizomes. Bot J Linn Soc 103: 127–132.

Perumalla, C.J., Peterson, C.A., and Enstone, D.E. (1990b). A survey of angiosperm species to detect hypodermal Casparian bands. I. Roots with a uniseriate hypodermis and epidermis. Bot J Linn Soc 103: 93–112.

Peterson, C.A. (1989). Significance of the Exodermis in Root Function. In Structural and Functional Aspects of Transport in Roots: Third International Symposium on ‘Structure and Function of Roots’ Nitra, Czechoslovakia, 3–7 August 1987, B.C. Loughamn, O. Gašparíková, and J. Kolek, eds, Developments in Plant and Soil Sciences. (Springer Netherlands: Dordrecht), pp. 35–40.

Peterson, C.A. and Perumalla, C.J. (1990). A survey of angiosperm species to detect hypodermal Casparian bands. II. Roots with a multiseriate hypodermis or epidermis. Bot J Linn Soc 103: 113–125.

Pfaffl, M.W. (2001). A new mathematical model for relative quantification in real-time RT-PCR. Nucleic Acids Res. 29: e45.

Pongrac, P., Baltrenaite, E., Vavpetič, P., Kelemen, M., Kladnik, A., Budič, B., Vogel-Mikuš, K., Regvar, M., Baltrenas, P., and Pelicon, P. (2019). Tissue-specific element profiles in Scots pine (*Pinus sylvestris L*.) needles. Trees 33: 91–101.

Ranathunge, K. and Schreiber, L. (2011). Water and solute permeabilities of Arabidopsis roots in relation to the amount and composition of aliphatic suberin. J Exp Bot 62: 1961–1974.

Robinson, N.J., Procter, C.M., Connolly, E.L., and Guerinot, M.L. (1999). A ferric-chelate reductase for iron uptake from soils. Nature 397: 694–697.

Ryan, C.G. (2000). Quantitative trace element imaging using PIXE and the nuclear microprobe. Int J Imaging Syst Technol 11: 219–230.

Ryan, C.G., Laird, J.S., Fisher, L.A., Kirkham, R., and Moorhead, G.F. (2015). Improved Dynamic Analysis method for quantitative PIXE and SXRF element imaging of complex materials. Nucl. Instrum. Methods Phys. Res. B: Beam Interact. Mater. At. 363: 42–47.

Sasaki, A., Yamaji, N., and Ma, J.F. (2016). Transporters involved in mineral nutrient uptake in rice. J Exp Bot 67: 3645–3653.

Schaaf, G., Schikora, A., Häberle, J., Vert, G., Ludewig, U., Briat, J.-F., Curie, C., and von Wirén, N. (2005). A putative function for the arabidopsis Fe-Phytosiderophore transporter homolog AtYSL2 in Fe and Zn homeostasis. Plant Cell Physiol 46: 762–774.

Serra, O., Soler, M., Hohn, C., Sauveplane, V., Pinot, F., Franke, R., Schreiber, L., Prat, S., Molinas, M., and Figueras, M. (2009). *CYP86A33*-targeted gene silencing in potato tuber alters suberin composition, distorts suberin lamellae, and impairs the periderm’s water barrier function. Plant Physiol 149: 1050–1060.

Shiono, K., Yoshikawa, M., Kreszies, T., Yamada, S., Hojo, Y., Matsuura, T., Mori, I.C., Schreiber, L., and Yoshioka, T. (2022). Abscisic acid is required for exodermal suberization to form a barrier to radial oxygen loss in the adventitious roots of rice (*Oryza sativa*). New Phytol 233: 655–669.

Shukla, V. and Barberon, M. (2021). Building and breaking of a barrier: Suberin plasticity and function in the endodermis. Curr Opin Plant Biol 64: 102153.

Singh, S.P., Vogel-Mikuš, K., Vavpetič, P., Jeromel, L., Pelicon, P., Kumar, J., and Tuli, R. (2014). Spatial X-ray fluorescence micro-imaging of minerals in grain tissues of wheat and related genotypes. Planta 240: 277–289.

Soukup, A., Armstrong, W., Schreiber, L., Franke, R., and Votrubová, O. (2007). Apoplastic barriers to radial oxygen loss and solute penetration: a chemical and functional comparison of the exodermis of two wetland species, *Phragmites australis* and *Glyceria maxima*. New Phytol 173: 264–278.

Soukup, A. and Tylová, E. (2018). Apoplastic Barriers: Their Structure and Function from a Historical Perspective. In Concepts in Cell Biology - History and Evolution, V.P. Sahi and F. Baluška, eds, Plant Cell Monographs. (Springer International Publishing: Cham), pp. 155–183.

Srivastava, A.K., Shankar, A., Nalini Chandran, A.K., Sharma, M., Jung, K.-H., Suprasanna, P., and Pandey, G.K. (2020). Emerging concepts of potassium homeostasis in plants. J Exp Bot 71: 608–619.

Ursache, R., Andersen, T.G., Marhavý, P., and Geldner, N. (2018). A protocol for combining fluorescent proteins with histological stains for diverse cell wall components. Plant J. 93: 399–412.

Vavpetič, P., Pelicon, P., Vogel-Mikuš, K., Grlj, N., Pongrac, P., Jeromel, L., Ogrinc, N., and Regvar, M. (2013). Micro-PIXE on thin plant tissue samples in frozen hydrated state: A novel addition to JSI nuclear microprobe. Methods Phys. Res. B: Beam Interact. Mater. At. 306: 140–143.

Vavpetič, P., Vogel-Mikuš, K., Jeromel, L., Ogrinc Potočnik, N., Pongrac, P., Drobne, D., Pipan Tkalec, Ž., Novak, S., Kos, M., Koren, Š., Regvar, M., and Pelicon, P. (2015). Elemental distribution and sample integrity comparison of freeze-dried and frozen-hydrated biological tissue samples with nuclear microprobe. Methods Phys. Res. B: Beam Interact. Mater. At. 348: 147–151.

Vogel-Mikuš, K., Pongrac, P., and Pelicon, P. (2014). Micro-PIXE elemental mapping for ionome studies of crop plants. Int. J. PIXE 24: 217–233.

Wang, P. et al. (2019). Surveillance of cell wall diffusion barrier integrity modulates water and solute transport in plants. Sci Rep 9: 4227.

Wang, P., Wang, C.-M., Gao, L., Cui, Y.-N., Yang, H.-L., de Silva, N.D.G., Ma, Q., Bao, A.-K., Flowers, T.J., Rowland, O., and Wang, S.-M. (2020). Aliphatic suberin confers salt tolerance to Arabidopsis by limiting Na+ influx, K+ efflux and water backflow. Plant Soil 448: 603–620.

Wu, C., Ye, Z., Li, H., Wu, S., Deng, D., Zhu, Y., and Wong, M. (2012). Do radial oxygen loss and external aeration affect iron plaque formation and arsenic accumulation and speciation in rice? J Exp Bot 63: 2961–2970.

Zeier, J. and Schreiber, L. (1997). Chemical Composition of hypodermal and endodermal cell walls and xylem vessels isolated from *Clivia miniata* (Identification of the biopolymers lignin and suberin). Plant Physiol 113: 1223–1231.

Zeier, J. and Schreiber, L. (1998). Comparative investigation of primary and tertiary endodermal cell walls isolated from the roots of five monocotyledoneous species: chemical composition in relation to fine structure. Planta 206: 349–361.

